# Fine Structural Features of Complex InDels and NHEJ Repair at Naturally Occurring Damage Sites in Normal Human Colon Crypts

**DOI:** 10.64898/2026.03.30.714948

**Authors:** Yong Hwee Eddie Loh, Michael R. Lieber, Timothy Okitsu, Cindy Okitsu, Jordan Wlodarczyk, Zarko Manojlovic, Chih-Lin Hsieh

**Affiliations:** Norris Medical Library, University of Southern California, Los Angeles, CA, USA; Department of Pathology, University of Southern California, Los Angeles, CA, USA; Department of Urology, University of Southern California, Los Angeles, CA, USA; Department of Surgery, University of Southern California, Los Angeles, CA, USA; current affiliation: Intermountain Health Saint Joseph Hospital, Denver, CO, USA

## Abstract

DNA repair in biochemical and genetic experimental systems permits a precise definition of enzyme requirements and mechanistic steps. Comparing these findings to repair events at naturally occurring damage sites in multicellular organisms is essential for confirming and expanding these insights into a physiologic context. However, heterogeneity in any normal cell population increases with each cell division, and the reliable detection of replication-independent DNA damage sites and their repair has been a major barrier. Here, we examine single human colon crypts, which harbor natural cell clones, using a novel whole-genome sequencing (WGS) method to identify complex insertion-deletion (indel) in the crypt stem cells. Analysis of complex indel events likely repaired by non-homologous end joining occurring in crypt stem cells permits inferences about the *in vivo* repair of naturally occurring DNA damage within physiologically-relevant chromatin in normal human cells.

## INTRODUCTION

Beyond simple point mutations, natural changes in genomic DNA can arise via several mechanisms. Replicative DNA synthesis can result in strand slippage events, which can give rise to insertions or deletions without DNA breakage, especially in repetitive sequence regions (1). Naturally arising DNA damage from background ionizing radiation and oxidative metabolism can lead to double-strand breaks (DSBs) in non-repetitive regions (2). There are four repair pathways for DSB in eukaryotic cells (3). The two main pathways are homologous recombination (HR) and nonhomologous end joining (NHEJ). HR occurs predominantly during S-phase, whereas NHEJ predominates during the remainder of the cell cycle (4). Single-strand annealing (SSA) and alternative end joining (aEJ) are two additional pathways that function primarily during DNA synthesis. HR and SSA use various lengths of strong sequence homology and can be readily distinguished from NHEJ (5, 6). The aEJ pathway is primarily involved in repairing collapsed replication forks and relies heavily on DNA polymerase theta (Pol Q) (7). The Pol Q transcript and protein levels are low in human colon epithelium, limiting the role of the aEJ pathway in repairing DNA damage in this tissue (8). In the human colon crypts, we can thus focus on the sites of replication-independent DNA damage, where DSB events are predominantly repaired by NHEJ.

In normal individuals, the components of the NHEJ machinery are constitutively expressed in all cells of the body at substantial levels (5, 9). The major components include Ku, DNA-PKcs, Artemis, polymerases mu and lambda, DNA ligase IV, XRCC4, and XLF (3). The mechanistic steps and roles of the components are discussed elsewhere (4). The structural biology underpinning the mechanistic steps has been elucidated (10). NHEJ is distinctive in that it does not require any DNA sequence homology or even microhomology (MH), and that entirely incompatible DNA ends can be joined. When microhomology is present, as is often pre-arranged in experimental systems, then the MH stabilizes the alignment of the two DNA ends in NHEJ, thereby favoring the outcome toward MH usage (4, 5). Polymerases mu and lambda are able to add nucleotides in either a template-independent or in a template-dependent manner (5, 11). Cellular studies of NHEJ have utilized various systems and approaches. Integrated target genes can be selected for event outcomes that permit drug selection of the target gene (12, 13). Additional insights have come from studies using transfected linear plasmids or DNA fragments followed by recovery (14, 15). Importantly, all these studies rely on cell lines in culture in which integration of target genes can be readily achieved.

Conventional whole genome sequencing (WGS) results typically reflect the consensus sequence of all the cells in a sample cell population. However, heterogeneity of the DNA sequence in a non-clonal cell population makes reliable detection of DNA damage sites repair extremely difficult, as discussed previously (1). For the first time, naturally arising somatic mutations can be comprehensively examined due to our technical breakthrough in generating high-depth and high-quality DNA sequences from natural cell clones, such as single human colon crypts, directly without DNA extraction (16, 17).

Based on the findings from biochemical and cellular studies, NHEJ events often appear to involve a small deletion with mis-incorporation of nucleotides; here, we term these complex indels. Curiously, almost no complex indel events were observed among the common small simple deletion and insertion (indels) variant calls from Mutect2 and Strelka, most likely because the variant-calling algorithms precluded the identification of the complex features. Using simulated complex indels, we determined that Freebayes is able to call complex indels in a single variant call much more often than some other variant callers (Y.H.E. Loh, in preparation). With our customized algorithm, we identified 3,041 potential complex indels from over 10^9^ Freebayes variant calls in our dataset of 106 single colon crypts. After removing variants shared by multiple individuals, 896 unique variants are manually examined on Integrated Genome Viewer (IGV) to confirm and validate, yielding 385 NHEJ events. The algorithm, the repair signature analysis, and the proposed repair mechanism are presented here.

## MATERIALS AND METHODS

### Tissue collection, crypt isolation, sequencing library construction, and whole genome sequencing

Collection of tissue from normal portions of colorectal samples, crypt isolation, and sequencing library construction have been described in detail previously (16, 17). A total of 106 crypt libraries and 21 bulk libraries are constructed from 21 individuals of age 10 months to 90 years old. After quality control, the sequencing libraries are pooled and sequenced on seven S4 flow cells using NovaSeq 6000 (Illumina, San Diego) S4 300 cycles reagent kit (v1.5) in the Keck Genomics Platform (KGP) Core facility at USC.

### Variant calling

The sequencing reads are assessed and processed using a validated in-house pipeline (based on GATK v4.2 best practices) from the KGP and the DRAGEN Somatic v 4.2.4 pipeline (Illumina Inc) as described previously (16) (17). PCR duplicate reads were identified and excluded from downstream variant-calling steps. Two germline callers, HaplotypeCaller and Freebayes (v1.2.0), and two somatic variant callers, Mutect2 and Strelka (v2.9.0), are used in the KGP pipeline.

### Algorithm for Identifying Potential NHEJ Events

NHEJ events frequently involve both nucleotide deletion and insertion (complex indel) at the same DNA damage site. SNVs and small indels (<50 bases) can be reliably identified by most somatic variant callers, such as Mutect2 and Strelka. We determined that events with NHEJ features are not observed among the indel variants called by either Mutect2 or Strelka. To reach this conclusion, we developed a test dataset comprising 2000 simulated mutations, including both complex and simple indel events across the chromosome 22 sequence of the human reference genome GRCh38. We used this dataset to test whether variant callers can identify each complex indel event in a single variant call. This test revealed that Freebayes can identify complex indels in single variant calls much more frequently than Mutect2, Strelka, and Haplotypecaller (Y.H.E. Loh, in preparation). Also, the DRAGEN pipeline does not perform as well as Freebayes in calling a complex indel in a single variant call. We then developed an algorithm to capture the NHEJ events from the variants identified by Freebayes (Fig. 1) as described briefly below.

**Figure 1.**
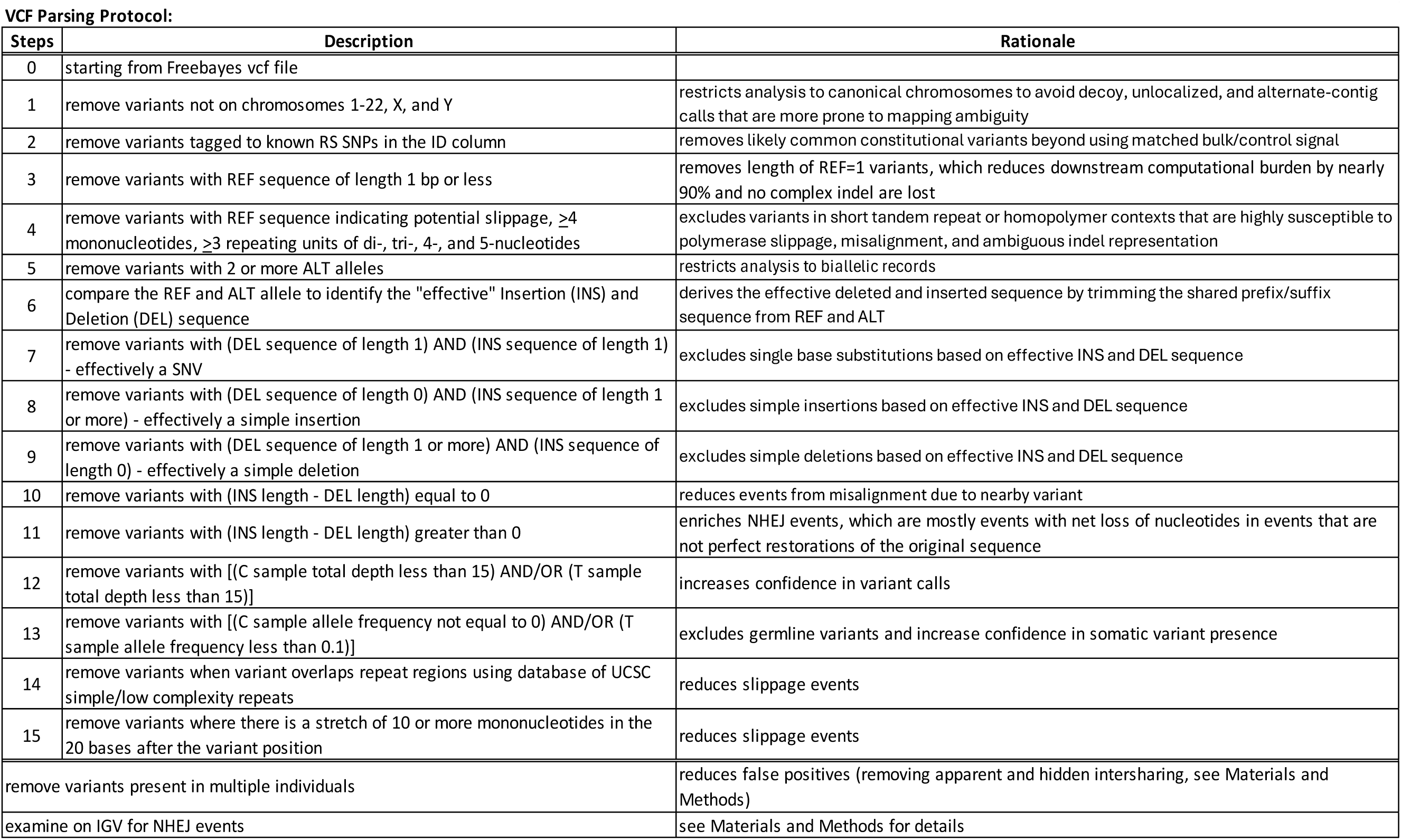
Workflow and rationale for identifying complex indel events from FreeBayes VCF.

Only variants on autosomes, and chromosomes X and Y from the Freebayes vcf are included in the current analysis. After removing known RS SNPs, which are likely to be common germline variants, variants with REF length ≤1 were removed to reduce downstream computational burden (Step #3 in Fig.1). This step removes nearly 90% of the reported variants, primarily single base substitutions that were reported with REF length = 1, and does not remove any complex indels. Variants with the following features are removed in the subsequent steps: sequence change due to slippage event (a stretch of ≥4 mononucleotides or a stretch of ≥3 repeating di-, tri-. 4-, or 5-nucleotide), multi-allelic variants, SNV, simple insertion, simple deletion, alternate allele frequency not equal to 0 in the bulk control (alternate allele present in the bulk control), alternate allele frequency <0.1 in the crypt, overlap with known repetitive regions, and variants with 10 or more mononucleotides in the 20 bases immediately downstream. Finally, variant calls appearing in multiple individuals are excluded, and variant calls appearing in multiple crypts of the same individual are collapsed into a single entry. Using our algorithm, a total of 3041 unique variants remained after processing more than 10^9^ total variants from 106 crypts with 21 matching bulk DNA controls.

Importantly, in our previous study analyzing simple insertions from this dataset, we found that hidden inter-sharing of variants across crypts and individuals is quite common (1). Sequencing reads of the alternate allele can be present in a crypt, but do not meet the criteria for a variant call due to read quality. Therefore, a variant may be called only in a single crypt while the alternate allele sequencing reads are present in multiple individuals, leading to hidden inter-sharing of the variant. We have developed a script previously to identify variants shared by more than a single individual (https://github.com/twewyttst777/VariantDuplicateCheck; (1). After removing hidden inter-sharing variants shared by more than a single individual from the unique variants, these variants are further manually examined on IGV to identify potential NHEJ events.

### Manual Inspection of Potential NHEJ Events

Events identified through the algorithm described above (n=896) are inspected manually on IGV for the following features: (a) length of deletion; (b) adjacent non-templated nucleotide insertions; (c) nearby misincorporations; (d) absence of repetitive sequences; (e) presence of potential base pairing of microhomology (MH); (f) at least two good quality alternate allele reads (quality score MAPQ ≥20); and (g) presence in both the KGP alignment and the DRAGEN alignment. Variants showed discernible NHEJ features (deletions, nucleotide insertions at or near the deletion boundary, and nearby nucleotide substitutions) supported by multiple good map quality sequencing reads are further analyzed for DNA repair processes.

Descriptive designation of the sequence alteration as presented on IGV is recorded for each variant. Sequence alterations can be interpreted differently depending on local sequence alignment. We note that the substitution inferred from IGV may not truly reflect the correct position of nucleotide change when we tabulate the nucleotide changes. The length of the DNA repair zone can only be inferred from the apparent DNA changes even though the actual zone of damage may be larger. We choose to define the minimum repair zone (MRZ) as the shortest possible zone from which the mutant allele can be derived. The number of nucleotides matching the reference genome between the altered nucleotides allows the determination of a potential mechanism for nucleotide additions. The potential MH usage is also considered by examining the sequences immediately upstream and downstream of each deletion, although the actual MH usage cannot be determined unequivocally.

## RESULTS

### General Features of the NHEJ Events Analyzed

From the 3041 algorithm-selected unique variants in 106 crypts from 21 individuals of various ages, 896 events are not shared by multiple individuals. These 896 events are examined manually on IGV to verify the total sequencing read count, alternate allele read count, map quality of the alternate allele sequencing reads, and the presence of the alternate allele in the alignment from the DRAGEN pipeline (Illumina, Inc.). The following features are further examined manually: (a) the length of deletion; (b) adjacent non-templated nucleotide insertions; (c) nearby misincorporations (non-templated single base changes); (d) sequences that could have potentially led to DNA slippage; and (e) the presence of potential MH for base pairing at the repair junction. Among the 896 events examined on IGV, 385 events show typical features of NHEJ (Supplementary Table S1), having nucleotide insertions or substitutions near the deletion and clearly without DNA slippage.

The minimum repair zone (MRZ) of each NHEJ event is determined by including the shortest region surrounding the start position of the variant demarcated by sequence alterations, including deletion, insertion, and base substitution sites, occurring consistently on the alternate allele (Supplementary Table S1). A base change could be an independent mutation if it is not present in all the alternate allele sequencing reads that harbor the deletion event a few nucleotides away. In contrast, it would strongly support that multiple changes occur as a single event if all alternate allele sequencing reads carry the same base substitution and the deletion. Therefore, the MRZ only includes non-germline alterations consistently occurring on the alternate allele chromosome within 50 nucleotides from the called position of the event.

Among the 385 NHEJ events, the event count decreases with the length of the MRZ as expected (Fig. 2). The longest MRZ observed is 43 nucleotides in length. Sixty percent (n=232) of the 385 events have an MRZ length of 2 or 3 nucleotides, and the MRZ is rarely (2.1%) more than 20 nucleotides in length (Figs. 2A and 2B). Interestingly, the MRZ length distribution does not vary substantially across the age distribution, in contrast to the number of events (Fig. 2C and section below).

**Figure 2.**
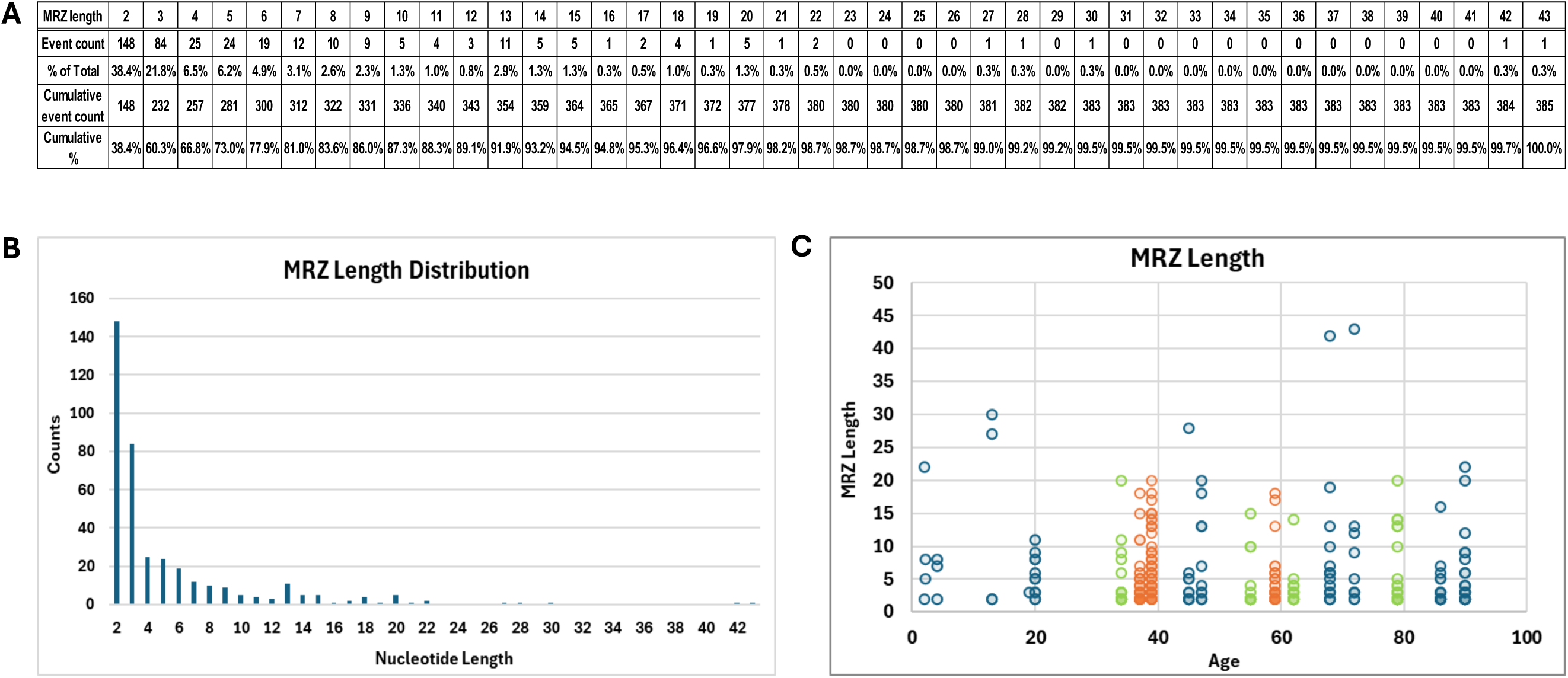
Minimum repair zone (MRZ) length distribution. MRZ is the smallest possible region of sequence alteration that can give rise to the alternate allele of the variant as described. (A) The length of MRZ from a total of 385 variants with NHEJ features are presented. (B) The distribution of MRZ length in 385 NHEJ events is displayed. (C) The distribution of MRZ length is displayed by age of the individuals. The color scheme of the dots are as described in Figure 3. The color fill of the dots is 90% transparent to allow multiple variants with the same MRZ to be visible (higher number of variants with the same MRZ have more saturated fill color).

### DNA damage accumulates with age, and chemotherapy/radiation combination treatment increases DNA damage in normal colon crypts

The number of NHEJ events showed a slight increasing trend with age (Fig. 3A and 3B). The three patients treated with a chemotherapy and radiation combination treatment show a much higher number of NHEJ events in their normal colon crypts (Fig. 3B, orange dots). When the three individuals with chemotherapy/radiation combination treatment are excluded from the analysis, the increase in NHEJ events observed in the remaining 18 individuals correlated with age much better, with R-squared value increasing from 0.137 to 0.712 (Fig. 3A and 3B orange shaded, and 3C). When only the 14 individuals with neither chemotherapy alone nor chemotherapy/radiation combination treatment (i.e., no treatment) are included in the analysis, the R-squared value increases to 0.855 (Fig. 3A and 3B, green shaded, and 3D). These findings indicate that our observation of NHEJ event increases can be interpreted as DNA damage accumulating with age at a rate of nearly 0.19 events per year in normal colon crypts (Fig. 3D).

**Figure 3.**
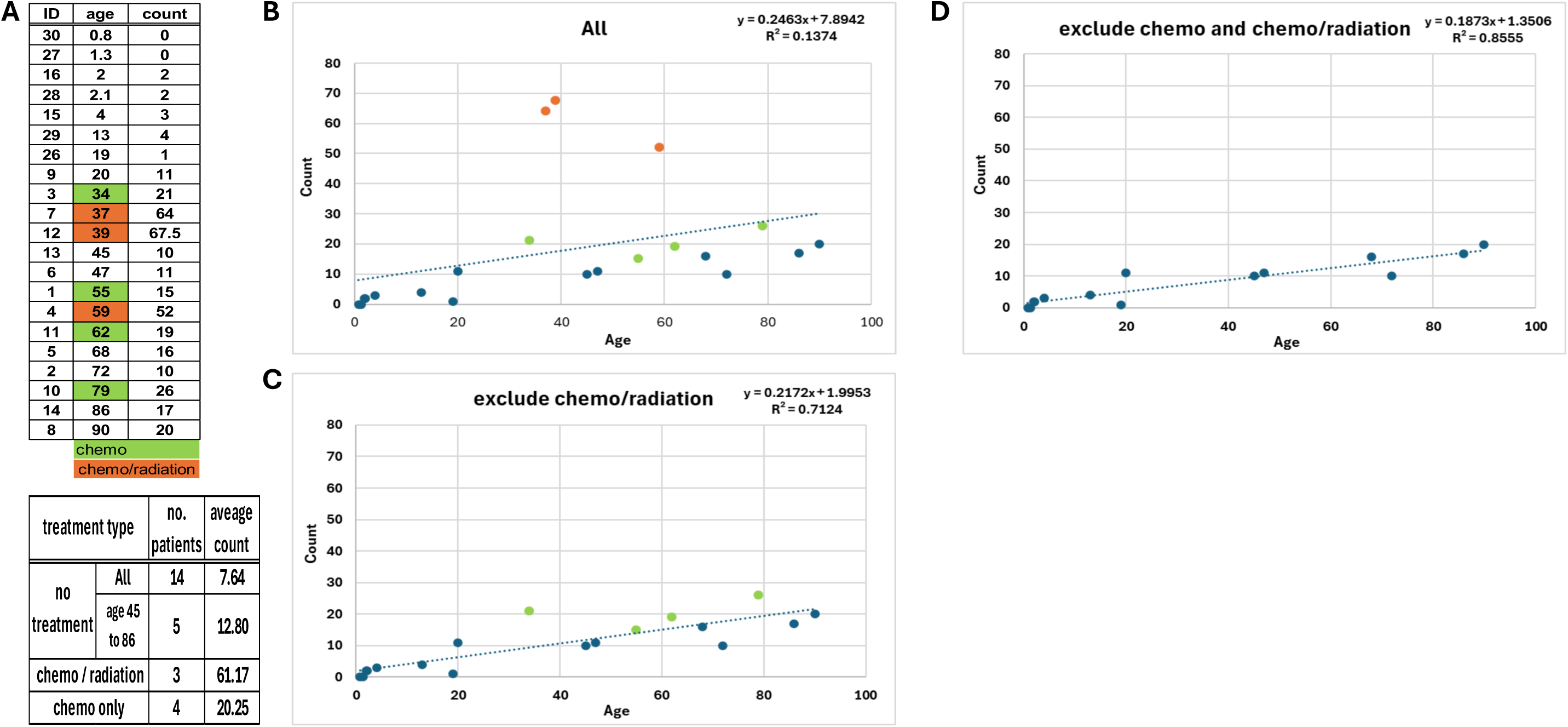
NHEJ event counts increase with age and medical treatment. (A) The total number of NHEJ events identified in each individual is presented with the average of three groups of patients calculated. Individual 0012 has 6 crypts instead of 5 like all the other samples; therefore, the total count for 0012 is adjusted by a factor of 0.833 (5/6). The burnt orange dots are patients with chemo/radiation combination treatment, the green dots are patients with chemo alone, and the blue dots are patients with no treatment. (B) The NHEJ event counts in all 21 individuals is graphed by age. (C) The NHEJ event counts in individuals without the chemo/radiation combination treatment (as marked in burnt orange in A) graphed by age. (D) The NHEJ event counts in individuals without treatment graphed by age. A linear trend line is displayed with the equation and the R-squared value in C and D.

Patients treated with chemotherapy alone have < 2-fold higher NHEJ event count (age 34 to 79 years old; average count 20.25, Fig. 3A) than patients of similar ages with no treatment (age 45 to 86 years old; average count 12.80). Since NHEJ event counts vary with age, it is appropriate to compare chemotherapy treated patients to patients of similar age range without treatment rather than compared to the patients of all ages without treatment (age 0.8 to 90 years old; average NHEJ event count = 7.64, Fig. 3A). Our observation from the small number of samples suggests that chemotherapy and radiation combination treatment (61.17 average mutation count) can potentially increase DNA damage three-fold over chemotherapy treatment alone (20.25 average mutation count) while chemotherapy treatment alone may increase DNA damage less than two-fold over no treatment in the normal colon crypts (12.80 average count) (Fig. 3A). In summary, chemotherapy and radiation combination treatment may potentially increase DNA damage burden about 5-fold over no treatment in the normal colon crypts (61.17 average mutation counts vs 12.80 average mutation counts).

### Nucleotide addition at colon NHEJ repair sites are mostly non-templated

Detailed information, including the reference sequence, alternate (mutant) allele sequence, the descriptive call, the final call, the MRZ, and the proposed repair mechanism of the 385 NHEJ events are summarized in Supplementary Table S1. Ten representative events are presented and described below to illustrate the complexity of interpretation and the final call of the alteration (Fig. 4).

**Figure 4.**
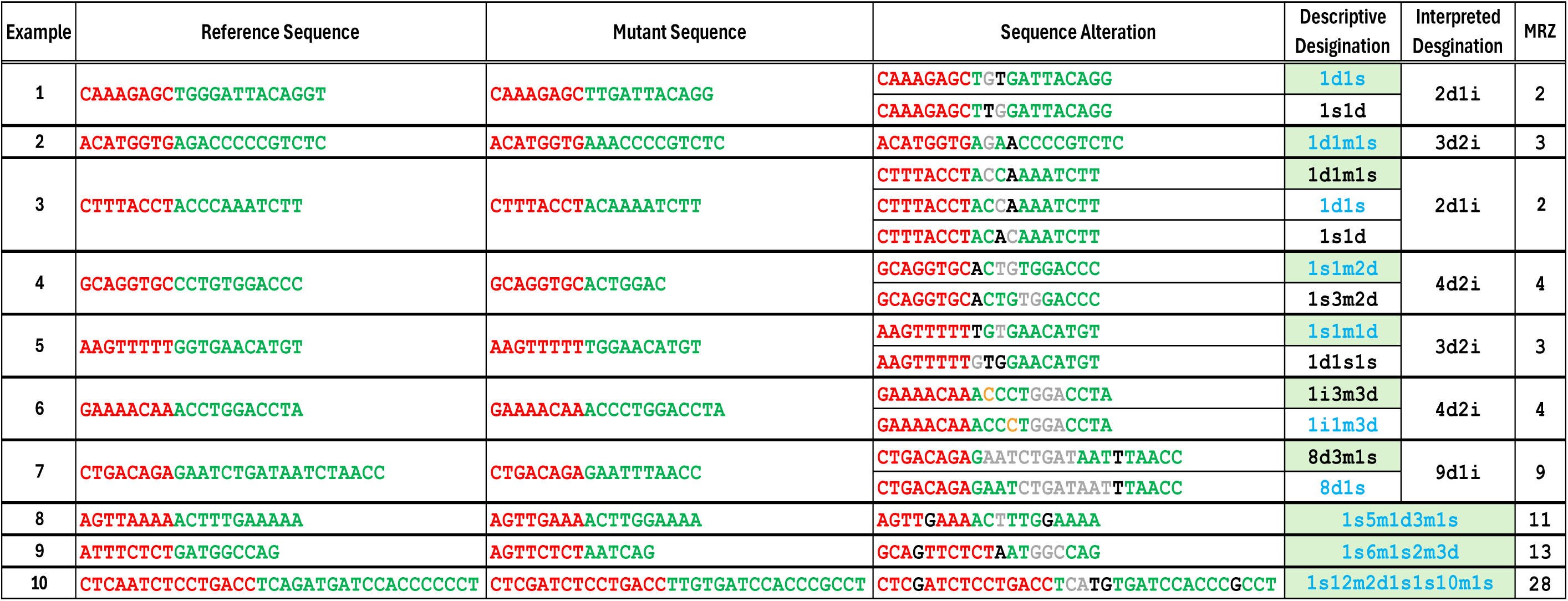
Sequence and interpretation of complex indel examples. The reference sequence is the nucleotide sequence in the reference genome GRCh38 flanking variant start position called by Freebayes. The mutant sequence is the sequence of the alternate allele. The sequence alteration illustrates the possible changes the alternate allele sequence can incur to result in the mutant sequence. The descriptive designation describes the observed changes with different possible local sequence alignments, and the alignment displayed on IGV is shaded with light green. The interpreted designation describes the zone of the shortest possible sequence change (MRZ) with the consideration of the most likely mechanism for nucleotide insertion. When the descriptive designation is the same as the interpreted designation, only a single entry is shown. The **red** letters represent nucleotides upstream and the **green** letters represent nucleotides downstream of the variant start position called by Freebayes. The **light grey** letters represent the deleted nucleotides. The **black** letters indicates the base changes. The **orange** letters represent the inserted nucleotides. The descriptive designations in **blue** are the ones with the shortest sequence changes.

As mentioned previously, DSBs in human colon epithelium are expected to be repaired predominantly by NHEJ. After DNA breakage occurs, polymerases mu and lambda can add nucleotides in either a template-independent or a template-dependent manner. Other alternatives to the local sequence alignment displayed in IGV are often present, and the interpretation of any sequence change can differ. We use the local sequence alignment that yields the shortest MRZ length to interpret DNA changes when multiple alignments are possible (Fig. 4, examples 3, 4, 6, and 7, blue letters in the Descriptive Designation column). The IGV alignment is used when there is no alternative alignment (Fig. 4, examples 2, and 8-10, Descriptive Designation column) or when the MRZ is the same for all alignments (Fig. 4, examples 1 and 5, blue letters in Descriptive Designation column).

The changes within the MRZ in the alternate allele frequently have zero or only a single nucleotide that match the germline sequence (Fig. 4, examples 1-7). These DNA changes are more likely to be the outcome of repair by polymerases adding random nucleotide(s) in a template-independent manner at the breakage site. Considering the mechanism involved, the interpretation of these NHEJ events is that a deletion of at least the size of the MRZ occurs at a DNA damage site followed by template-independent addition of nucleotides by DNA polymerases at the DNA ends before end joining (Fig. 4, Interpreted Designation column). It is important to note that the position of base substitution can be ambiguous in these events with non-templated nucleotide addition (Fig. 4, examples 1, 3, and 5). In contrast, when the MRZ in the alternate allele harbors more than a single nucleotide matching the germline sequence, the likelihood of template dependence of nucleotide addition increases (Fig. 4, examples 8-10). The position of base substitutions present in the templated addition events are typically clearly assignable.

Based on the potential mechanism of nucleotide addition of these NHEJ events as described above, we classified the NHEJ events observed into three categories: templated addition, non-templated addition, and exception (Fig. 5). Among the 385 events, 42 (10.9%) involve templated addition of nucleotides and 333 (86.5%) have non-templated nucleotide addition after DNA breakage and local end resection. The remaining ten (2.6%) events of exception appear to involve more complex processing of the broken DNA ends, and assessment of template involvement is unclear.

**Figure 5.**
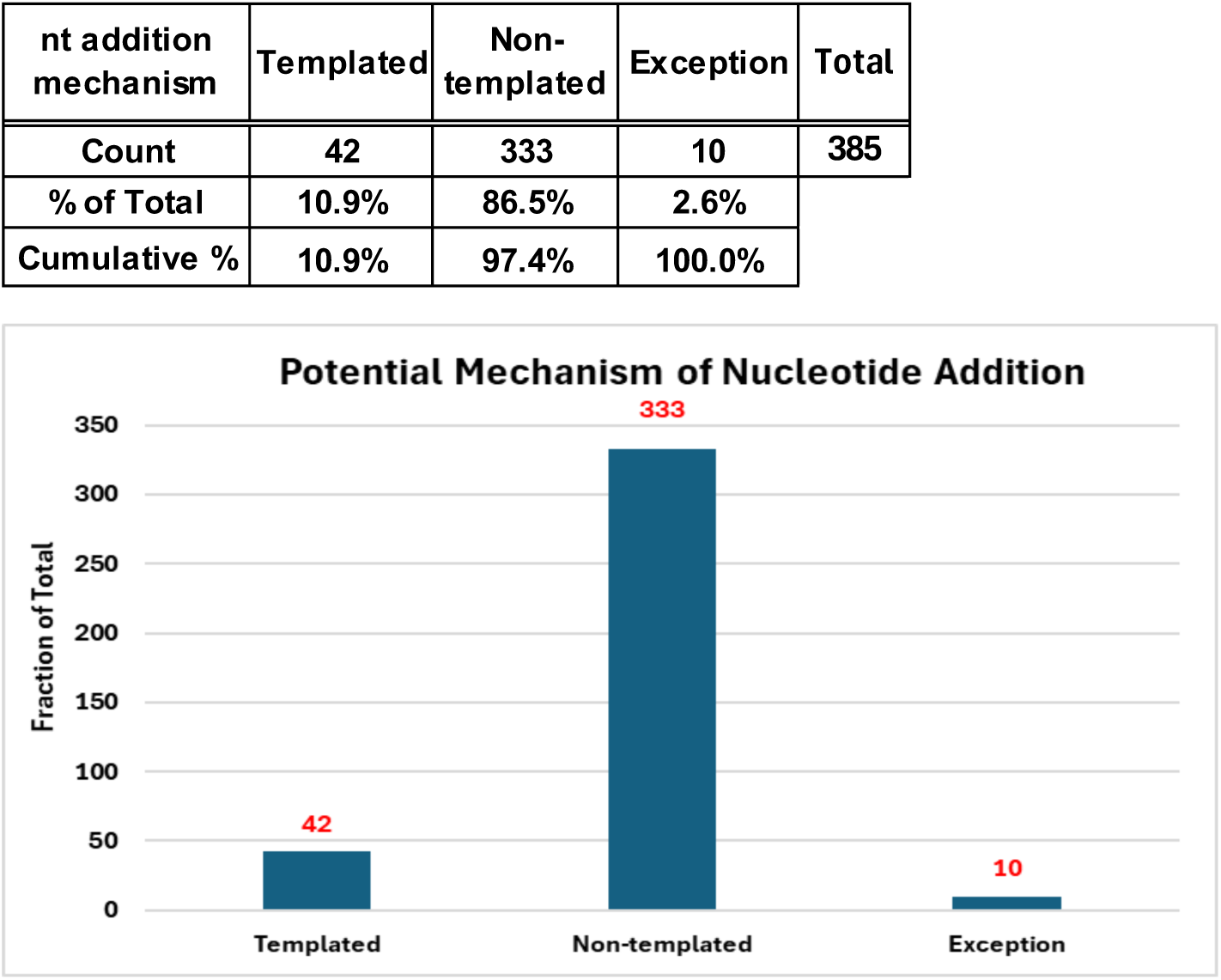
Distribution of NHEJ events by repair mechanism. The nucleotide addition can be classified as templated addition and non-templated addition. Some events may involve more than a single mechanism or being influenced by the adjacent sequences and classified as exception.

### The base content of nucleotide insertion is biased toward A and T for non-templated insertion

Despite the lack of certainty on the exact location of the nucleotide insertion as mentioned earlier, the content of the inserted nucleotide is much more reliable. We analyzed the 486 total nucleotides added in the 385 NHEJ events (Fig. 6). The ratio of A and T (A/T) to C and G (C/G) nucleotides are similar in the events repaired by templated addition of nucleotides (A/T=47.9%, C/G=52.1%) and the exception events repaired by a more complex mechanism (A/T=42.1%, C/G=57.9%). Interestingly, the non-templated nucleotide additions are 72.8% A/T and 27.2% C/G. Our findings indicate an A/T preference for the non-templated additions (see Discussion).

**Figure 6.**
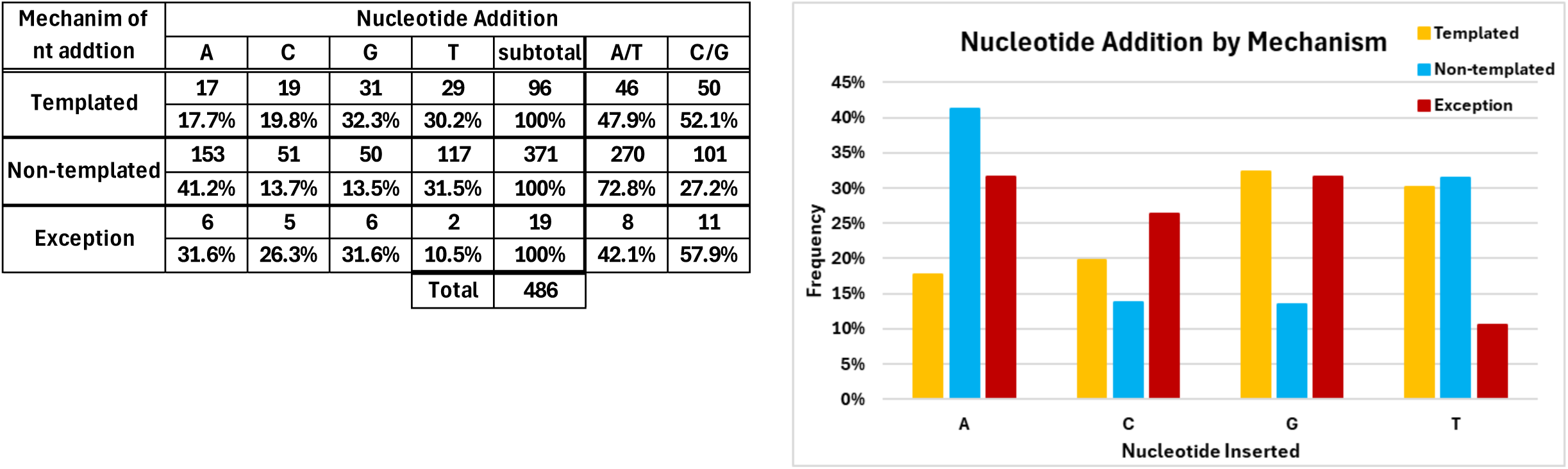
Content of nucleotide by repair mechanism. Nucleotides inserted by different mechanisms, templated, non-templated, and exception, are tabulated and graphed.

### Microhomology usage is uncommon at the NHEJ repair sites

We assessed for any MH in the 385 NHEJ events, though MH is not required in NHEJ. No potential MH usage is observed in 289 of the 385 NHEJ events (75.1% of the total, Fig. 7). The longest microhomology observed is 3 nucleotides. The likelihood of random match of the nucleotides by chance for one to three bases if all four nucleotides are present in equal frequency is somewhat higher than the observed MH candidates (Fig. 7A). Therefore, the apparent 2 or 3 bases of possible MH may be the result of chance matching and not represent actual MH usage. The absence of reliance on MH is consistent with NHEJ events observed *in vivo* (4). As in experimental systems, polymerase mu or polymerase lambda addition that matches the one or two nucleotides in a 3’ overhang on the other DNA end of the DSB would not be discernible as MH usage and has appropriately been referred to as occult microhomology (18).

**Figure 7.**
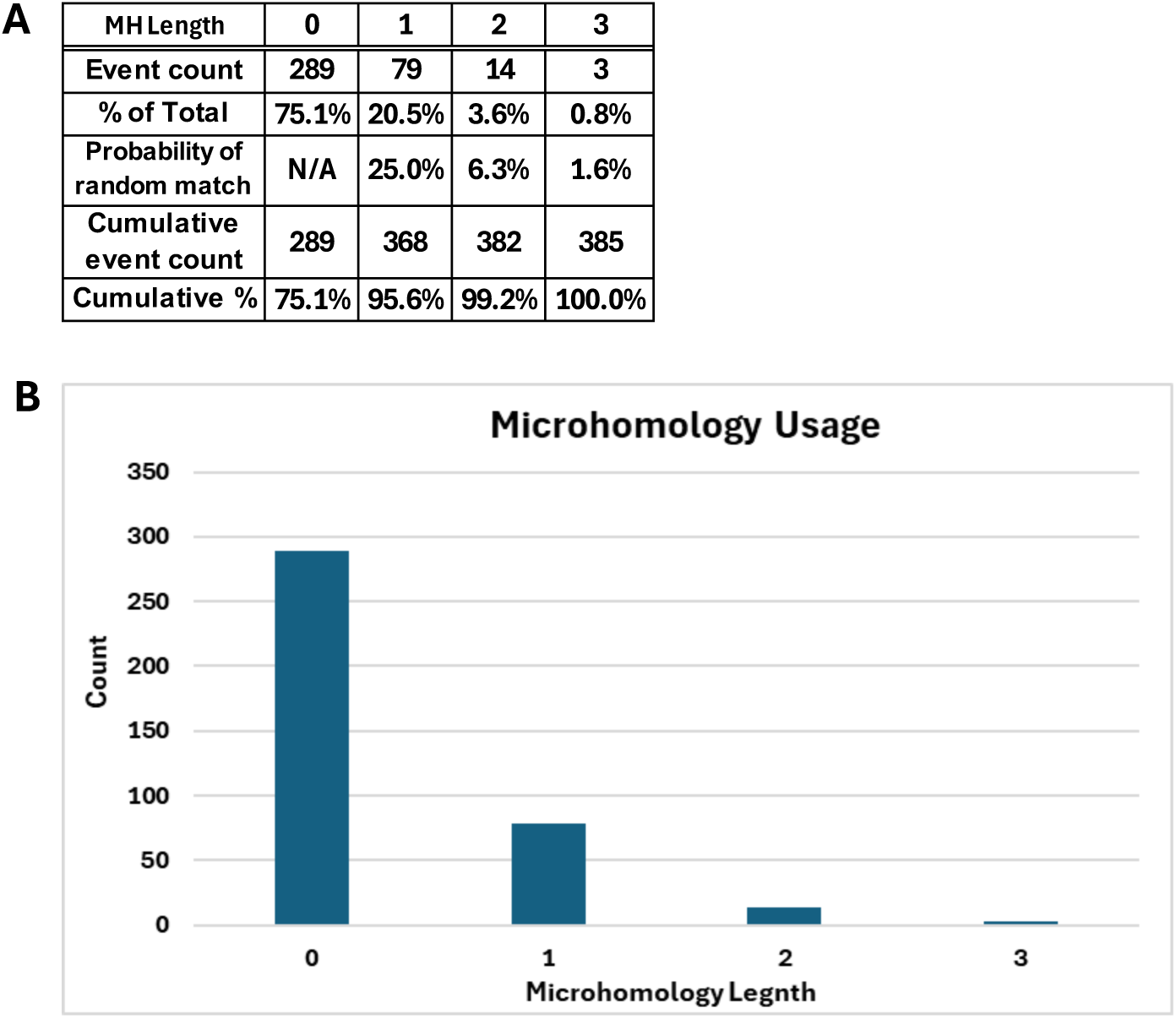
Microhomology usage in end joining. Nucleotide sequences at the beginning and end of each NHEJ event are compared with the flanking sequences to identify potential microhomology usage. (A) Tabulation of microhomology usage. (B) The count of different number of bases of microhomology usage is graphed.

## DISCUSSION

Unlike in immortalized and neoplastic cells where both NHEJ and some level of alternative end joining (aEJ) are found, mammalian somatic cell genomic DSB events are mostly repaired by NHEJ in primary cells (5, 6). Particularly in human colon epithelium, where Pol Q (the polymerase for aEJ) expression is low, NHEJ is nearly the only mechanism for DSB repair in the absence of obvious long homology usage to support HR or SSA. In general, NHEJ repair events include nucleotide insertions and/or substitutions at the breakage site in addition to the deletion generated by end processing after DNA breakage (i.e., complex indels). When deletion and insertion are equal in length, the inserted nucleotide(s) can be 1) a complete match to the original sequence without any trace of breakage or 2) can have mismatch(s) to the original sequence and appear as one or more base substitutions at the repair site. No DNA change will be detected in the first situation, and the repair signature cannot be distinguished from SNV events in the second situation. When the insertion is longer than the deletion at the repair site, the event cannot be easily distinguished from a simple insertion (1). When the insertion is shorter than the deletion upon repair, the repair signature can either 1) appear to be a simple deletion if the added nucleotides are indistinguishable from the original sequence or 2) have clearly identifiable nucleotide additions within the longer deletion that represents potential NHEJ events.

In our analysis of simple insertions, only 77 events of the 4,972 unique insertions from 106 single colon crypts are clearly DNA breakage events (1). Only seven of the 77 insertions with DNA breakage are complex indels as confirmed by manual validation of BAM files in IGV. We have determined that most variant callers are unable to identify the vast majority of complex indels as single variants. We tested several commonly used variant callers using simulated complex indel events and identified Freebayes as the best for calling a complex indel in a single variant call instead of multiple variant calls (Y.H.E. Loh, in preparation). We also determined that NHEJ events are best captured by identifying variants with more nucleotides deleted than inserted at a specific chromosomal location, as discussed above. We then developed an algorithm to identify potential NHEJ events among the variant calls from Freebayes (Fig. 1). The candidate NHEJ events were manually examined in IGV. It is important to note that our approach underestimates the total number of complex indels for reasons described above. In addition, Freebayes does not identify all complex indels, and our algorithm requires that the deletion size exceed the insertion size. Therefore, NHEJ events that do not result in any overall deletion, such as those with only base changes or those with larger insertions, are not included in our analysis. Among the seven complex indels found in our insertion analysis of the same dataset, some are likely NHEJ events in which the insertion is longer than the deletion (1). In contrast, a total of 385 NHEJ events with the insertion shorter than the deletion are identified from Freebayes vcf using our algorithm, indicating Freebayes outperforms Mutect and Strelka for complex indel identification.

Our study here of NHEJ in primary human somatic cells shows an increase in NHEJ repair lesions with age among individuals without therapeutic radiation or chemotherapy treatment. We found that therapeutic radiation substantially increases NHEJ events beyond the slight increase with chemotherapy treatment alone. These increases are not due to any known germline pathological mutations in the DNA damage response and repair loci in these individuals, as examined in an independent analysis (Z. Manojlovic, in preparation). It has been difficult in previous studies to assess the impact of ionizing radiation on DSB repair sites in human primary cells by previous biochemical or cellular transfection experiments. Our findings on the DSB impact in human primary cells distant from treated colon cancer neoplasms, yet within the radiation field, are very relevant to the understanding of radiation effects on normal cells (19, 20).

Our study of complex indels and NHEJ repair has the advantage of analyzing these events in their natural nuclear chromatin environment on a genome-wide scale. In biochemical or cellular transfection studies, genetic selection and specific naked DNA end structure may artificially constrain or affect the repair events observed. Such approaches often involve generating two DNA ends at separate points in time, minutes or more apart, raising the possibility that the two DNA ends are processed independently before being joined. Also, experimental systems using purified proteins or crude extracts assume that repair reflects the natural nuclear environment. Transfections of non-replicating DNA may occur in the cytoplasm or other unintended cellular compartments. Therefore, our quantified genomic observations in the unaltered human body within normal cells as presented here provide an essential and critical dataset.

Ionizing radiation is known to cause both single- and double-strand DNA damage, typically with one DSB for every 20 to 40 SSB (3). The distribution and nature of damage of the DSB in the native nuclear context of primary human cells has been unclear. Here we find that the number of complex indels repaired in a manner consistent with NHEJ is substantially elevated in individuals receiving chemotherapy and radiation combination treatment. Since we find that chemotherapy alone only has a minimal impact here on the number of NHEJ events in the normal colon crypts of the treated patients, we propose that the substantial increase of NHEJ events in patients with chemotherapy and radiation combination treatment is primarily due to the radiation. Our findings here closely parallel those in a previous study reporting increased single base substitutions (SBSs) in colon organoids established from chemotherapy treated colon cancer samples (21). Although not specifically noted in that previous study, our analysis of their data indicates that radiation also appears to exaggerate SBS burden beyond that of chemotherapy alone, and these SBS are mostly the result of single-strand DNA damage.

Interestingly, the MRZ size of the NHEJ events in patients with chemotherapy and radiation combined treatment or only chemotherapy treatment is similar to that of patients without treatment. This observation indicates that while ionizing radiation increases the number of DNA breakage events, it does not impact the repair process itself to any detectable level. This simple observation also suggests an important aspect of the trajectory of the ionizing radiation damage through chromatin. Given the tight wrap of DNA around the histone core particles, the ionizing radiation particle track could have been expected to cause DSB sites ∼83 bp apart, which is the distance between points of the two duplexes wrapped within one nucleosome. However, the MRZ, which reflects the size of the deletion and insertion in each event, does not appear to differ between treated and untreated patients. While the interpretations of this observation are complex, one possibility is that ionizing radiation particles generate a local distribution of damage that is not substantially different from the free radicals of normal oxidative metabolism that cause many naturally arising DSB (2). Most of the NHEJ events in our study have an MRZ length of 5 nucleotides or less (∼73%), suggesting that the breaks on the two strands of the DNA duplex are within this small zone, regardless of the cause of the DNA damage in most cases. The limited MRZ size also suggests that the strand break sites leading to a DSB are generally within 4 or 5 nucleotides of one another.

Our analysis of the content of nucleotide additions at DNA breakage sites reveals the preference of A/T over C/G when the nucleotide is added by the non-templated mechanism. Preferential non-templated addition of A has also been observed *in vivo* and *in vitro* in a wide range of organisms by many DNA polymerases (as summarized in (22))(23-26). Our *in vivo* human genomic observations here indicate a preference of adenine addition by polymerases mu and polymerase lambda, which are the enzymes involved in the NHEJ events, and *in vitro* studies are consistent with this (27, 28).

As mentioned earlier, NHEJ is distinctive in that it does not require sequence homology or microhomology for end joining. We observe no microhomology usage in a high percentage of the 385 events (75.1%). The actual frequency of end joining without microhomology can be higher because the apparent one to three bases of homology can occur by random chance (Fig. 7A). These findings are consistent with what has been observed *in vitro*, but it is valuable to have this comparison with normal primary somatic cells incurring DSB within the physiologic environment of the cell nucleus within the human body (4, 5).

The limitation of our approach is that we do not know the DNA break site as precisely as in experimental biochemical systems. Thus, the MRZ is inferred as the shortest DNA region in which a combination of deletions, insertions, and substitutions can give rise to the alternate (mutant) allele. The most likely mechanism for broken DNA processing and repair can be inferred from the features of the observed changes. Our observation and interpretation of the naturally occurring NHEJ events provide a novel perspective that adds to the rich set of already-existing data obtained using experimental systems.

## ACKNOWLEDGEMENT

We would like to thank the Norris Comprehensive Cancer Center Translational Pathology Core for the sample collection. We would like to acknowledge the Norris Comprehensive Cancer Center Molecular Genomics Core and the Keck Genomics Platform at the University of Southern California for the sequencing work. This work is supported by funds from NIA R01 AG 067615 and the Catherine and Joseph Aresty Endowment to CLH and ZM. MRL was supported by NIGMS R35118009.

